# REDUCED CACNA1C EXPRESSION PRODUCES ANHEDONIC REACTIONS TO PALATABLE SUCROSE IN RATS: NO INTERACTIONS WITH JUVENILE OR ADULT STRESS

**DOI:** 10.1101/2024.08.21.608991

**Authors:** Patricia Gasalla, Kerrie Thomas, Lawrence Wilkinson, Jeremy Hall, Dominic Dwyer

**Author notes:** Address for correspondence: Patricia Gasalla Canto School of Psychology Cardiff University Tower building, 70 Park Place, Cardiff, CF10 3AT. UK.

## Abstract

Genetic variation in *CACNA1C*, which encodes the alpha-1 subunit of Cav1.2 L-type voltage-gated calcium channels, is strongly linked to risk for psychiatric disorders including schizophrenia, bipolar disorder, and major depression. Here we investigated the impact of mutations of one copy of *Cacna1c* on rats’ hedonic responses to palatable sucrose (assessed using the analysis of consumption microstructure). In addition, we also investigated the effects of combining either juvenile or adult stress with the manipulation of *Cacna1c*. Across three experiments *Cacna1c*^+/-^ rats displayed attenuated hedonic reactions to sucrose compared to WT littermate controls, despite the *Cacna1c*^+/-^ rats retaining sensitivity to sucrose concentration in terms of the amount of consumption. Unexpectedly, juvenile stress enhanced rather than reduced hedonic reactions to sucrose, while adult stress did not have clear hedonic effects. The effects of *Cacna1c* manipulation did not interact with either juvenile or adult stress. The fact that *Cacna1c*^+/-^ rats display a clear analogue of anhedonia – a reduction in the positive hedonic reactions normally elicited by highly palatable sucrose – a symptom observed transdiagnostically across psychiatric disorders linked to *CACNA1C*, suggests this model may play a valuable role in the translational fiinvestigation of anhedonia.

## 1- Introduction

Schizophrenia (SCZ) and Bipolar Disorder (BD), which can both exhibit psychotic features, are severe adult psychiatric disorders affecting approximately 3% of the population (Grande, Berk, Birmaher, & Vieta, 2016; McDaid & Park, 2022; Owen, Sawa, & Mortensen, 2016). Both are highly heritable (Cardno et al., 1999; Sullivan, Kendler, & Neale, 2003) with a large degree of overlap of genetic risk between them (as well as between both and major depression) (Lee et al., 2013). Although the psychotic symptoms (e.g., delusions, hallucinations) of SCZ are relatively amenable to pharmacological treatment, the negative symptoms (e.g. anhedonia, amotivation) are not, and these are strongly associated with functional impairment and remain poorly understood mechanistically (Correll & Schooler, 2020; Owen et al., 2016). Similarly, reasonable acute pharmacological treatments for mania in BD exist but with risk of exacerbating depressive symptoms, and antidepressants can risk exacerbating mania (Grande et al., 2016). Thus, there is a clear unmet need for understanding the biological basis for the negative symptom of psychosis.

Genomic studies of SCZ and BD have demonstrated the importance of genetic variation in voltage gated calcium channels (VGCCs) in risk for these conditions (Mullins et al., 2021; Trubetskoy et al., 2022). VGCCs play a critical role in regulating calcium influx and synaptic plasticity in the central nervous system (Berger & Bartsch, 2014). Single nucleotide polymorphism (SNP) variants in *CACNA1C*, which encodes the pore-containing α1 subunit of the L-Type VGCC Cav1.2, has strong and replicated genome-wide association with both SCZ (Ripke et al., 2014; Trubetskoy et al., 2022) and BD (Ferreira et al., 2008; Mullins et al., 2021). Moreover, altered dosage of *CACNA1C* is important for risk, with evidence suggesting a particular role for low dosage in limbic regions (Gershon et al., 2014; Jaffe et al., 2020; Roussos et al., 2014) but see also (Bigos et al., 2010). In light of the transdiagnostic display of symptoms such as anhedonia, it is notable that genetic variation in VGCCs is also associated with risk for MDD (Bhat et al., 2012; Rao et al., 2016).

The investigation of *CACNA1C* contributions to the risk for neuropsychiatric disease has been furthered by the development of a rat model which reflects altered dosage seen in SCZ and BD (Jaffe et al., 2020). Prior characterisation of these *Cacna1c*^+/-^ rats (with 4bp deletion in exon 6) confirms reduced *Cacna1c* mRNA and Cav1.2 α1 subunit protein expression between 17% and 48% depending on brain region (A. L. Moon, Brydges, Wilkinson, Hall, & Thomas, 2020; Sykes et al., 2019). These rats also display aberrant Ca^2+^ signalling and downregulation of the ERK pathway in hippocampus (Tigaret et al., 2021).

Activation of the ERK pathway with a BDNF mimetic rescued synaptic plasticity and attenuated latent inhibition deficits characteristic of cognitive dysfunction observed in psychosis (Tigaret et al., 2021). Such results confirm the heterozygous *Cacna1c*^+/-^ rat model appropriately reflects biological dysfunctions related to genetic variation in *CACNA1C* and demonstrates a pathway whereby genetic variation in *CACNA1C* can disrupt synaptic plasticity processes, which in turn contribute to cognitive deficits associated with neuropsychiatric disorders. However, such cognitive deficits are only a subset of the relevant symptoms and reward processing deficits central to the negative symptom cluster observed transdiagnostically remain under-investigated.

Importantly, the negative symptoms are not monolithic, reflecting separable dysfunctions in reward processing: including direct hedonic deficits (anhedonia), motivational problems (amotivation), and reward-related cognitive biases - with variation in presentation of symptoms across individuals. Here, we investigate a rodent analogue of anhedonia, or the reduction in positive hedonic reactions typically elicited by positive stimulation, by examining the microstructure of consumption behaviour. Rodents typically produce clusters of licks separated by pauses, and the mean number of licks per cluster displays a positive monotonic relationship with the concentration of palatable sucrose solution (e.g., J. D. Davis & Perez, 1993; J. D. Davis & G. P. Smith, 1992; Spector, Klumpp, & Kaplan, 1998), a negative relationship with unpalatable solution such as quinine (Hsiao & Fan, 1993; Spector & St John, 1998), as well as being sensitive to pharmacological interventions known to affect hedonic reactions in humans (Asin, Davis, & Bednarz, 1992; Higgs & Cooper, 1998). Critically, lick cluster size is not simply a proxy for consumption: studies of conditioned taste aversion and preference have also shown that palatability and consumption can dissociate (e.g., D. M. Dwyer, 2009b; D. M. Dwyer, Boakes, & Hayward, 2008; D. M. Dwyer, Burgess, & Honey, 2012; D. M. Dwyer, Gasalla, & Lopez, 2013; D. M. Dwyer, Pincham, Thein, & Harris, 2009). In the present experiments we used the analysis of lick cluster size to provide a means of selectively assessing hedonic responses. In this light, a reduction in lick cluster size compared to control animals while consuming highly palatable sucrose solution is a clear analogue of anhedonic reactions (Clarkson, Dwyer, Flecknell, Leach, & Rowe, 2018; Wright, Gilmour, & Dwyer, 2020).

In addition to genetics, environmental risk factors are also known to play an important role in the development of psychotic illnesses (Marangoni, Hernandez, & Faedda, 2016; Owen et al., 2016). Stress in both adulthood (Agid, Kohn, & Lerer, 2000) and childhood (Varese et al., 2012) is reliably associated with psychiatric disease in humans. Moreover, variation in *CACNA1C* is associated with levels of perceived stress in humans (Pennington et al., 2020), while reduced expression of *Cacna1c* in mouse Nucleus Accumbens resulted in increased susceptibility to social stress (Terrillion, Francis, Puche, Lobo, & Gould, 2017), and the relationships between stress susceptibility and *Cacna1c* may vary with age (Dedic et al., 2018). Importantly, rats exposed to short variable stressors pre-pubertally have long lasting alterations in behaviour and brain activation including altered learning and increased anxiety (N. M. Brydges, Hall, Nicolson, Holmes, & Hall, 2012; Nichola Marie Brydges et al., 2018; N. M. Brydges, Wood, Holmes, & Hall, 2014; Jacobson-Pick, Audet, Nathoo, & Anisman, 2011), with the same pre-pubertal stress rat model producing a decrease in the expression of *Cacna1c* which is sustained into adulthood (A. Moon, Brydges, Thomas, & Hall, 2019). These results suggest that genetic and environmental risk factors for psychosis may converge on VGCCs, through altered expression of *CACNA1C*. However, neither the impact of this juvenile (or adult) stress procedure on hedonic behaviour, nor its potential interaction with direct manipulation of *Cacna1c* has been investigated. Therefore, in addition to examining the effect of low-dose cacna1c on hedonic behaviour using the heterozygous *Cacna1c*^+/-^ rat model (Experiment 1), Experiment 2 combined this manipulation with the effects of juvenile stress, while Experiment 3 examined the effects of the same stress protocol delivered in adult *Cacna1c*^+/-^ rats.

## 2- Experiment 1

### 2.1. Method

#### 2.1.1. Subjects

Ninety *Cacna1c* hemizygous (*Cacna1c*^+/−^) rats (HET) on a Sprague Dawley background (TGR16930, Horizon, Sage Research Labs, USA) and wild-type (WT) littermates were bred at Cardiff University, UK (for further details of the model see - A. L. Moon et al., 2020; Sykes et al., 2019). Animals were and housed in single-sex and mixed-genotype groups of 2–3 in standard cages (38cm × 56cm × 22cm) under 12hr/12hr light/dark cycle. The housing conditions included poplar bedding (Datesand, UK) and enrichment items in the home cage. For enrichment, each cage was equipped with a rat tunnel (125 x 90 x 5 mm) and aspen wood bricks (10 x 2 x 2 cm, rt chew sticks) (Datesand, UK). The weight range for females was 194 to 251 grams, and for males it was 328 to 488 grams at the beginning of the experiment. Four days before the start of the experiment, rats were moved to a food deprivation schedule with food daily access to maintain animals between 85 and 90% of their *ad lib* weights. Animals had access to ad lib water throughout the experimental sessions. All experimental manipulations took place during the light phase of the cycle. We used both female and male WT and HET rats: 21 HET female, 18 WT female, 29 HET male, and 22 WT male at approximately 10 weeks old. All experiments were conducted in accordance with local ethics guidelines, the UK Animals (Scientific Procedures) Act 1986 (PPL P0EA855DA held by Jeremy Hall) and the European Communities Council Directive (1986/609/EEC).

#### 2.1.2. Stimulus and apparatus

This experiment used 4%, 8%, and 16% (w/w) sucrose solutions made with deionized water. Training and testing phases took place in a room containing 6 identical conditioning boxes (38 x 24 x 21 cm: Height x Width x Depth; Med Associates). The side walls of the boxes were constructed from aluminium, whereas the front, back, and the ceiling were made from clear acrylic. The floor was formed from 19 steel rods (4.8 mm diameter, 16 mm apart) placed above a stainless-steel tray. One aluminium wall contained two 1 cm diameter holes, one at the left and one at the right side, each 5 cm from the respective wall and from the floor of the box. These holes allowed for drinking bottles consisting of a steel spout and 50 mL bottle to be accessed by rats inside the box, while the bottles were in the forward position.

Bottles were automatically advanced/withdrawn at the beginning/end of each session. Licks were recorded by a PC running MEDPC (Med Associates, St Albans), which measured contacts with a bottle spout to the nearest 0.01 s. Bottles were weighted with a scale accurate to 0.01g before and after each session. A weighing boat was placed below the hole and outside the box to collect any spill produced by retracting the bottle, ensuring accuracy in the consumption measure.

#### 2.1.3. Procedure

At the beginning of the experiment, animals were moved to a food restriction schedule until they reached 90% of ad libitum body weight. Training started four days after being moved to food restriction. The training phase consisted of 10 sessions in which animals received 10 minutes of access to 8% (w/w) sucrose solution daily in order to habituate them to the boxes in the experimental room, timing and procedures. Lick cluster size and consumption were measured for each animal in each session. Drinking sessions took place between approximately 9 a.m. and 2 p.m. every day (with the order counterbalanced across HET/WT and male/female animals) and rats received a measured food ration in their home cages in the afternoon. Once all animals displayed stable consumption and licking patterns, they were moved to test phase. Half of the animals were presented with 4% (w/w) sucrose solution for 10 minutes daily for four consecutive days, while the other half were presented with 16% sucrose solution. Animals were then presented with the alternative solution across another four daily sessions. This means that the animals that previously received 4% sucrose solution were administered the 16% sucrose solution, and vice versa.

#### 2.1.4. Data analysis

Total consumption and mean lick cluster size were the main dependent variables. Lick cluster size was defined as a group of licks separated by intervals of less than 0.5 seconds, a criterion that has been extensively employed in our laboratory (D. M. Dwyer, 2008, 2009a; D. M. Dwyer, Figueroa, Gasalla, & Lopez, 2018; Dominic M Dwyer, Gasalla, & López, 2019), following the original proposal by Davis (John D Davis, 1973, 1989; John D Davis & Gerard P Smith, 1992). For the tests, the average of the 4 sessions at each sucrose concentration was calculated for each animal. Mixed ANOVA was then performed with concentration (4% vs 16%) as within-subject variable, and sex (male vs female) and genotype (HET vs WT) as between-subject factors. All test reported here used a criterion for significance of *p* = .05.

### 2.2. Results

Figure 1 shows the consumption (1A) and lick cluster size (1B) data over test sessions. Mixed ANOVA performed on the consumption data during test revealed significant main effects of solution concentration, *F*(1,86) = 114.47, *p* < .001, MSE = 2.44, ƞ^2^ = .57, and sex, *F*(1,86) = 7.62, *p* = .007, MSE = 12.02, ƞ^2^ = .08. However, the analysis revealed no significant effect of genotype, *F*(1,86) = 1.70, *p* = .196, MSE = 12.02, ƞ^2^ = .01, nor any significant interaction between factors (largest F for concentration by sex by genotype interaction, *F*(1,86) = 2.21, *p* = .141, MSE = 2.44, ƞ^2^ = .02). While males consumed more sucrose than females, there was no effect of genotype, and both HET and WT animals had similar consumption levels. The fact that consumption of 16% sucrose was higher than that of 4% sucrose for both genotypes indicates both HET and WT animals were sensitive to sucrose concentration and showed a preference for higher sucrose concentrations.

**Figure 1.**
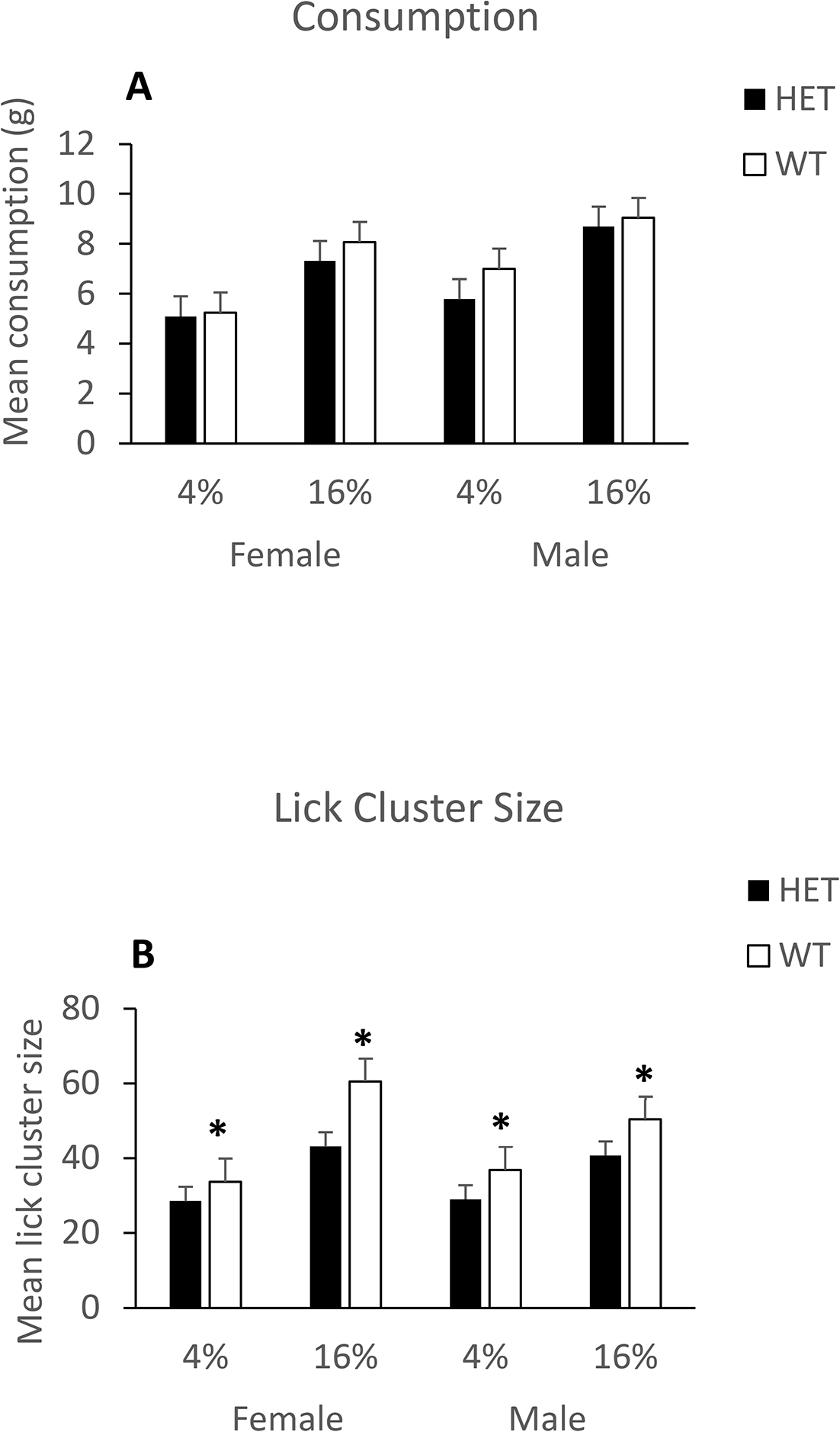
Average test data (over 4 sessions) for female and male HET and WT animals. Test duration was 10 minutes per day and animals had ad lib access to an either 4% sucrose solution or 16% sucrose solution on different days. (A) Mean sucrose consumption in grams, and (B) mean lick cluster size displayed by animals. Error bars represent the standard errors of the mean (SEM), * indicates significant pairwise comparisons between genotypes (HET vs. WT) with p < .05.

The same analysis performed on the palatability data revealed significant main effects of solution concentration, *F*(1,86) = 52.71, *p* < .001, MSE = 229.83, ƞ^2^ = .38, and importantly, of genotype, *F*(1,86) = 7.10, *p* = .009, MSE = 616.78, ƞ^2^ = .08. There was no significant effect of sex, *F*(1,86) = 0.35, *p* = .554, MSE = 616.78, ƞ^2^ = .01, nor any significant interaction between factors (largest F for concentration by sex interaction, *F*(1,86) = 3.13, *p* =.080, MSE = 229.83, ƞ^2^ = .03). These results showed that HET animals had a lower lick cluster size than WT, regardless of solution concentration and sex. Thus, there was clearly an analogue of anhedonia in the *cacna1c* HET animals that cannot be explained as a failure to detect differences in sucrose concentration given that HET animals did show concentration effects in the amount of sucrose consumed.

However, before considering the implications of this result, Experiment 2 will examine the potential interaction between cacna1c manipulation and juvenile stress, and Experiment 3 will examine potential interaction between cacna1c manipulation and adult stress. In particular, in Experiment 2 we exposed female and male *Cacna1c* HET & WT rats to three mild stressors from postnatal days 25-27: forced swimming, an elevated platform, and electric shocks. In adulthood, we tested their response to palatable sucrose as in Experiment 1. Given the fact that rats with reduced *Cacna1c* expression displayed attenuated hedonic reactions to sucrose in Experiment 1, and juvenile stress can produce long-lasting reductions in *Cacna1c* expression, we would expect both direct reduction of *Cacna1c* in the HET rats and juvenile stress to result in lower lick cluster sizes when consuming sucrose as adults.

## 3- Experiment 2

### 3.1. Method

#### 3.1.1. Subjects, stimulus and apparatus

One hundred and twenty-four *Cacna1c*^+/−^ rats (HET) and wild-type (WT) littermates were bred at Cardiff University for Experiment 2. The experiment was performed with two similar cohorts of animals: in the Juvenile Stress (JS) groups there were 13 HET females (4/9 in cohort one and two respectively), 18 HET males (12/6), 17 WT females (12/5), and 19 WT males (9/10); in the Control groups (non-stressed littermates that were just weighted and handled on same days as the stress animals) there were 17 HET females (11/6), 18 HET males (10/8), 11 WT females (4/7), and 11 WT males (9/2). The weight range for females was 180 to 268 grams, and for males it was 222 to 306 grams at the beginning of the experiment. Housing conditions, food restriction schedule, testing equipment, and sucrose solutions were the same as in Experiment 1.

#### 3.1.2. Early-life stressors

Between postnatal days 25 and 27, half of the animals underwent a short-term mild stress protocol, adapted from, and previously used in our laboratory (N. M. Brydges et al., 2012; Nichola Marie Brydges et al., 2018; N. M. Brydges et al., 2014; Jacobson-Pick et al., 2011). The animals were moved to an experimental room different from the holding room and sucrose test room. Sessions started at 10 am and finished by approximately 3 pm. On postnatal day 25, animals were subject to 10 minute forced swim session inside an opaque swimming tank measuring 25 cm in height and 34 cm in diameter. The tank, with a capacity of 12 litters contained approximately 6 litters of water at a consistent temperature of 25 ± 1 °C. Animals were released facing the wall of the tank and retrieved after 10 minutes. Immediately after retrieval, the rats were dried with a towel and closely monitored for 30 minutes in the experimental room. Animals were further monitored in the home cage and did not show prolonged signs of distress following removal from the water. The following day (postnatal day 26), animals from stress groups were exposed to an elevated platform (15 x 15 cm^2^ , 115 cm high) for three 30-minutes sessions separated by 60 minutes each round. Finally, on postnatal day 27, animals were exposed to 6 × 0.5 second shocks (0.5 mA) separated by 30 second intervals. Shocks were delivered in a room containing 4 identical conditioning boxes (30 x 24 x 21 cm: Height x Width x Depth; Med Associates) housed within individual sound and light attenuating chamber. The side walls of the boxes were constructed from aluminium whereas the front, back, and the ceiling were made from clear acrylic. The floor was formed from 19 steel rods (4.8 mm diameter, 16 mm apart) placed above a stainless-steel tray. Illumination was provided by standard house light (40 mA) mounted in back wall of each chamber. The electric shock was delivered using a scrambled shocker (Campden Instruments Ltd. Model HSCK1000). The rats were video monitored over the sessions.

Animals were then returned to the home cage and remained undisturbed, apart from regular welfare checking and general husbandry until reaching adulthood at postnatal day 70. All animals caged together belong to the same group (JS or Control) to avoid any potential stress transfer between animals.

#### 3.1.3. Behavioral Procedure and data analysis

The behavioural testing procedures were the same as in Experiment 1 with the exception that animals had access to 6 hours 8% sucrose solution before training in their home cages to reduce neophobic effect and promote consumption in the experimental boxes. Drinking sessions took place in the morning (approximately 10 a.m. to 3 p.m.) and rats received a measured food ration in their home cages in the afternoon. As in Experiment 1, total consumption and mean lick cluster size were used as the main dependant variables. The average of the four test sessions at each sucrose concentration was calculated for each animal and then subjected to mixed ANOVA with concentration (4% vs 16%) as a within-subject variable, and stress-group, sex, and genotype as between-subject factors.

### 3.2. Results

Figure 2A and 2B shows the mean sucrose consumption in grams for females and males respectively. Mixed ANOVA performed with the consumption data revealed significant main effects of solution concentration, *F*(1,116) = 360.36, *p* < .001, MSE = 5.55, ƞ^2^ = .76, and sex, *F*(1,116) = 5.82, *p* = .017, MSE = 19.53, ƞ^2^ = .05 However, the analysis revealed no significant main effects of genotype, *F*(1,116) = .43, *p* = .514, MSE =19.53, ƞ^2^ =.01, or stress, *F*(1,116) = 3.31, *p* = .071, MSE =19.53, ƞ^2^ = .03, nor any significant interaction between factors (largest F for concentration by sex by genotype by stress interaction, *F*(1,116) = 1.80, *p* = .183, MSE = 5.55, ƞ^2^ = .01). These results are consistent with Experiment 1. While males consumed more sucrose than females, there was no genotype or stress effect on the amount of sucrose consumed. Again, the higher consumption of 16% sucrose than 4% sucrose indicates that all groups were sensitive to sucrose concentration and showed a preference for higher concentrations.

**Figure 2.**
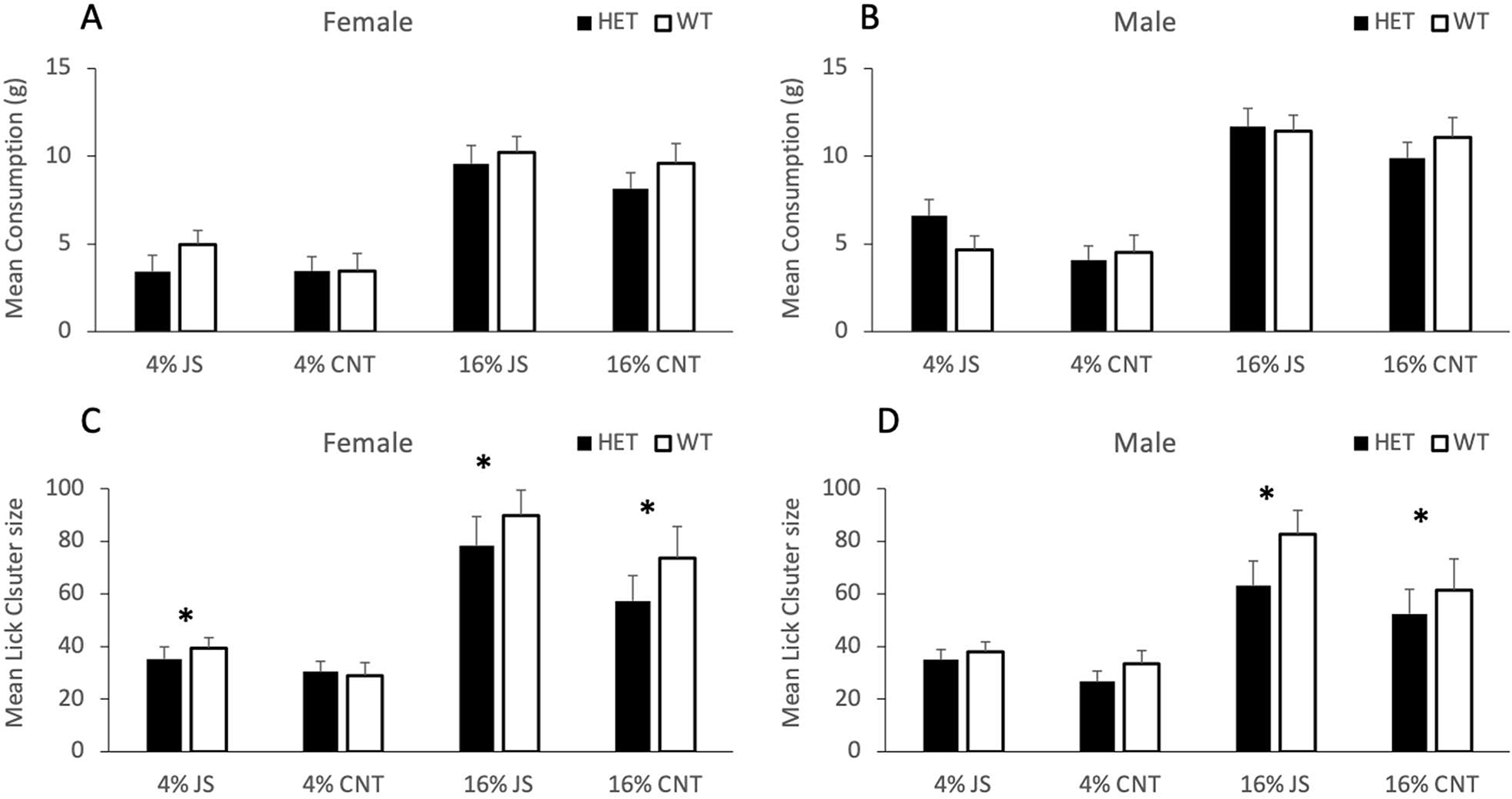
Average test data (over 4 sessions) for female and male HET and WT animals as a function of juvenile stress exposure (JS) vs control (CNT). Test duration was 10 minutes where animals had ad lib access to an either 4% sucrose solution or 16% sucrose solution on different days. (A and B) Mean sucrose consumption in grams for females and males respectively, and (C and D) mean lick cluster size displayed by female and male animals respectively. Error bars represent the standard errors of the mean (SEM), * indicates significant pairwise comparisons between genotypes (HET vs. WT) with p < .05.

Figure 2C and 2D shows the mean lick cluster size for females and males respectively. Analysis of the palatability data revealed significant main effects of solution concentration, *F*(1,116) = 104.47, *p* < .001, MSE = 754.47, ƞ^2^ = .47, and critically, of genotype, *F*(1,116) = 3.98, *p* = .048, MSE = 1092.13, ƞ^2^ = .03, as well as of stress-group *F*(1,116) = 8.03, *p* = .005, MSE = 1092.13, ƞ^2^ = .07. Animals generally display higher lick cluster sizes for the higher sucrose concentration, but more importantly HET animals displayed a lower lick cluster sizes compared to WT regardless of the sex of the animal and previous stress experience. In addition, rats that experienced juvenile stress displayed higher lick cluster size than controls animals suggesting higher palatability for sucrose. The remainder of the analysis revealed no significant effect of sex *F*(1,116) = 1.38, *p* = .242, MSE = 1092.13, ƞ^2^ = .01, nor any significant interaction between factors (largest F for concentration by genotype interaction, *F*(1,116) = 2.36, *p* = .127, MSE = 754.47, ƞ^2^ = .02).

Experiment 2 replicated the effects of direct *Cacna1c* manipulation seen in Experiment 1, namely that HET animals showed a clear reduction in hedonic reactions to sucrose compared to WT. However, while there was also a clear effect of stress-group, it was in the opposite direction to our expectations – rats subject to juvenile stress displayed higher hedonic reactions to sucrose as adults compared to controls. These two effects appeared to be independent of each other. We will return to the unexpected positive hedonic effects of juvenile stress in the general discussion, but first, Experiment 3 will examine adult stress.

Experiment 3 was performed generally as in Experiment 2, save that the stress protocol was performed when the rats were adults (i.e. approximately 8-9 weeks old).

## 4- Experiment 3

### 4.1. Method

#### 4.1.1. Subjects, stimulus, apparatus, and stressors

One hundred and three *Cacna1c*^+/−^ rats (HET) and wild-type (WT) littermates were bred at Cardiff University for Experiment 3. In the Adult Stress (AS) groups there were 13 HET females (6/7 animals in cohort one and two respectively, 13 (8/5) HET males, 12(5 /7) WT females and 15(9/6) WT males; in the Control groups (non-stressed littermates that were just weighted and handled on same days as the stress animals) there were 12 (6/6)HET females, 11 (7/4)HET males, 9 (5/4)WT females and 18(8/10) WT males. The weight range for females was 222 to 306 grams, and for males it was 327 to 505 grams at the beginning of the experiment. Housing conditions, food restriction schedule, testing equipment, and sucrose solutions were the same as in Experiment 2, with the exception that training lasted for eight sessions and test for six sessions because steady consumption levels were reached more rapidly than in previous experiments.

The stressors used were the same as in Experiment 2, save that they were delivered once animals reach adulthood (approximately 8-9 weeks old). Stress sessions started at 10 am and finished at approximately 3 pm. A week after delivering the last stressor the behavioural procedure began.

#### 4.1.2. Behavioral Procedure and data analysis

The behavioural testing procedures were the same as in Experiments 1 and 2. Drinking sessions took place in the morning (approximately 10 a.m. to 3 p.m.) and rats received a measured food ration in their home cages in the afternoon. Again, total consumption and mean lick cluster size were used as the main dependant variables. The average of the three sessions at each solution concentration was calculated for each animal and then subjected to mixed ANOVA with concentration (4% vs 16%) as a within-subject variable, and stress-group, sex, and genotype as between-subject factors. Nine animals (two WT stress males, two WT control males, two HET stress males, one WT stress female, one HET control male, and one HET control female) were excluded from this analysis either because there was no lick cluster data for one or both concentrations (either because consumption was minimal or the lick recording itself failed).

### 4.2. Results

Figure 3A and 3B shows the mean sucrose consumption in grams for females and males respectively. As in previous experiments, consumption was higher for 16% than 4% sucrose, and higher for males than females. In addition, the male/female difference in consumption appeared to be reduced in animals subject to adult stress. Mixed ANOVA performed with the consumption data revealed significant main effects of solution concentration, *F*(1,86) = 171.97, *p* < .001, MSE = 6.33, ƞ^2^ = .667, and sex, *F*(1,86) = 27.48, *p* < .001, MSE = 25.77, ƞ^2^ = .24, as well as a significant sex by concentration interaction, *F*(1,86) = 9.69, *p* = .003, MSE = 6.33, ƞ^2^ = .10 and a significant sex by concentration by genotype interaction, *F*(1,86) = 6.90, *p* = .010, MSE = 6.33, ƞ^2^ = .07. Follow up analysis of this 3-way interaction revealed that male HET animals consumed less 4% sucrose than their WT littermates, *F*(1,86) = 5.60, *p* = .020, MSE = 20.74, ƞ^2^ = .061, but there were no other HET/WT differences (largest *F*(1,86) = 0.88, *p* = .352, MSE = 11.36, ƞ^2^ = .01 for the comparison in female animals for 4% sucrose). However, the analysis revealed no main differences between genotype, *F*(1,86) = 0.31, *p* = .581, MSE = 25.77, ƞ^2^ < .01, stress-group , *F*(1,86) = 0.56, *p* = .457, MSE = 25.77, ƞ^2^ = .01, nor any other significant interaction between factors (largest F for stress-group by sex interaction, *F*(1,86) = 2.88, *p* = .093, MSE = 25.77, ƞ^2^ = .03).

**Figure 3.**
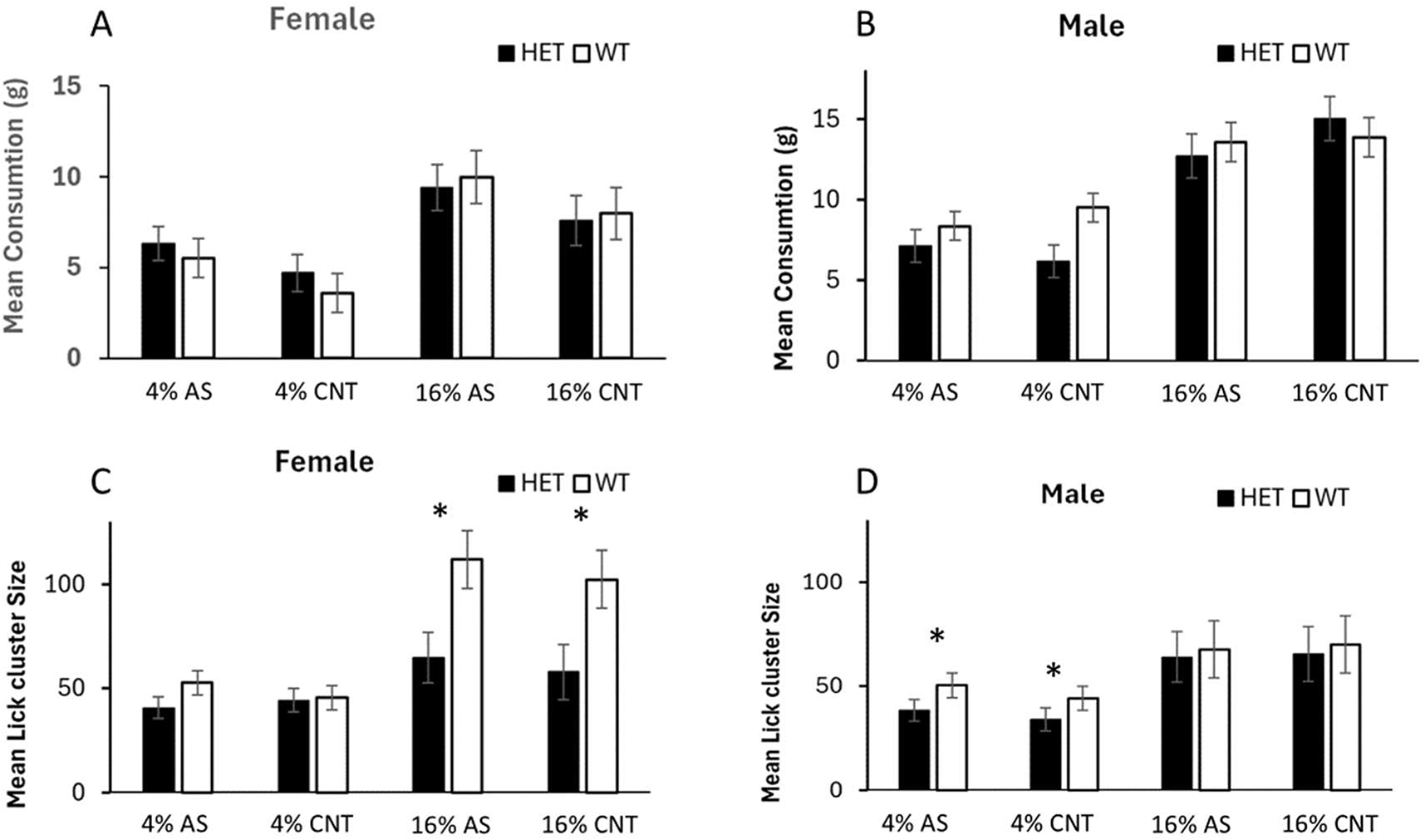
Average test data (over three sessions) for female and male HET and WT animals as a function of adult stress exposure (AS) vs control (CNT). Test duration was 10 minutes where animals had ad lib access to an either 4% sucrose solution or 16% sucrose solution on different days. (A and B) Mean sucrose consumption in grams for females and males respectively, and (C and D) mean lick cluster size displayed by female and male animals respectively. Error bars represent the standard errors of the mean (SEM), * indicates significant pairwise comparisons between genotypes (HET vs. WT) with p < .05.

Figure 3C and 3D shows the mean lick cluster size for females and males respectively. Contrary to juvenile stress, there was no suggestion that adult stress enhanced lick cluster size, however, we replicated the previously observed reduction in lick cluster size for cacna1c rats. ANOVA revealed a significant main effect of solution concentration, *F*(1,86) = 67.53, *p* < .001, MSE = 690.75, ƞ^2^ = .44, a main effect of genotype, *F*(1,86) =8.29, *p* = .005, MSE = 1588.78, ƞ^2^ = .09, and a significant genotype by concentration by sex interaction *F*(1,86) = 8.84, *p* = .004, MSE = 690.75, ƞ^2^ = .09. Follow-up analysis of this interaction revealed that there was a significant difference between HET and WT female animals for 16% sucrose, *F*(1,86) = 11.81, *p* < .001, MSE = 1932.70, ƞ^2^ = .12, but not for 4% sucrose, *F*(1,86) = 1.40, *p* = .241, MSE = 346.83, ƞ^2^ = .02, as well as a significant difference between HET and WT male animals for 4% sucrose, *F*(1,86) = 4.26, *p* = .042, MSE = 346.83, ƞ^2^ = .05, but not for 16% sucrose, *F*(1,86) = 0.11, *p* = .743, MSE = 1932.70, ƞ^2^ = .01. In addition, there were no significant effects of stress group, *F*(1,86) = 0.33, *p* = .566, MSE = 1588.78, ƞ^2^ < .01, or sex, *F*(1,36) = 3.37, *p* = .070, MSE = 1588.78, ƞ^2^ = .04, nor any other interaction (largest F for the concentration by sex interaction, *F*(1,86) = 3.00, *p* = .087, MSE = 690.75, ƞ^2^ = .03).

Thus, unlike in Experiment 2 with juvenile stress, there was no clear enhancement of palatability reactions following adult stress. In addition, the general pattern of lower hedonic reactions for the *Cacna1c* HET rats was again replicated here, albeit the genotype effect was more prominent for female rats at high sucrose concentration, but more prominent for male rats at low sucrose concentration.

## 5. General discussion

The most clear and consistent result across Experiments 1, 2, and 3 is that *Cacna1c*^+/-^ rats displayed lower lick cluster sizes when consuming sucrose than WT littermate rats, indicating a clear effect of *Cacna1c* genotype on hedonic responses. This was present in both male and female rats and did not interact with either juvenile or adult stress. Importantly, this effect on lick cluster size was present despite the fact that *Cacna1c*^+/-^ rats remained sensitive to differences in sucrose concentration in terms of its impact on the overall amount of sucrose consumed. Thus, the lick analysis suggests the presence of a true hedonic deficit that cannot be reduced to a failure to detect sucrose itself. In short, *Cacna1c*^+/-^ rats display a clear analogue of anhedonia – they show a reduction in the positive hedonic reactions normally elicited by highly palatable sucrose. In addition to this reliable genotype effect, juvenile stress unexpectedly resulted in an increase in hedonic reactions to sucrose (Experiment 2), while a similar effect was not observed after adult stress (Experiment 3). These results have implications for both the investigation of the biological mechanisms contributing to anhedonia and for the understanding of stress and resilience. We will cover each of these issues in turn.

The highly reliable observation of a defect in hedonic reactions to sucrose in *Cacna1c*^+/-^ rats is not only consistent with the fact that variation in this VGCC-encoding gene is associated with risk for multiple psychiatric disorders where anhedonia is a key symptom, but also consistent with the specifics impacts of variation in *CACNA1C* in humans on reward processing. For example, using a probabilistic reward-learning task Lancaster and colleagues (Lancaster, Heerey, Mantripragada, & Linden, 2014) found that individuals with the A risk allele carriers (AA/AG) of the *CACNA1C* rs1006737 genotype had a deficit in reward responsiveness compared with the non-risk (GG) genotype group. In addition, the amount of amygdala activation in response to a reward reversal learning task also differed between the A risk allele carriers (AA/AG) of rs1006737 and the non-risk (GG) genotype group (Wessa et al., 2010). Moreover, A risk allele carriers of rs1006737 also displayed altered resting-state functional connectivity across a network of brain regions, including those associated with emotion and reward processing (Jiang et al., 2023). Taken together with the current results, these studies of *CACNA1C* variation in humans suggest the presence of a translationally- preserved deficit in the behavioural responses to rewards and in the function of reward-related brain networks. Overall, these results support the view that genetic variation in *CACNA1C* may contribute to anhedonia trans-diagnostically across a range of psychiatric presentations.

Turning to the effects of stress, we had expected that stress would be likely to produce a reduction in hedonic responses to sucrose, and for this to interact with the effect of *Cacna1c* variation. These expectations were based specifically on the fact that the juvenile stress procedure we used has been shown to impact on anxiety and learning (N. M. Brydges et al., 2012; Nichola Marie Brydges et al., 2018), as well as producing a decrease in the expression of *Cacna1c* itself (A. Moon et al., 2019). More generally, across a range of species and potential stressors, exposure to stress is associated with depressive behaviours (Andersen, 2015). Despite these expectations, juvenile stress resulted in an increase in hedonic reactions to sucrose, while adult stress had no significant impact on hedonic response. Although counter to our expectations, such a result is not unprecedented - a recent meta-analytic review noted that exposure to developmental stressors had very heterogeneous effects including a number of reports of positive effects despite the overall estimate of the effect size being negative (Eyck, Buchanan, Crino, & Jessop, 2019). Indeed, the fact that stress may be associated with positive outcomes in some situations is a key contributor to theoretical analyses suggesting that that exposure to stress can protect against later challenges such as the inoculation-stress hypothesis (Levine, 1962; Lyons, Parker, & Schatzberg, 2010) or the related match/mismatch hypothesis (Hartmann & Schmidt, 2020). In this light, the higher hedonic reactions after juvenile stress (but not adult stress) could suggest that early negative experiences have produced some resilience to subsequent challenges. While intriguing, such a possibility remains speculative given that it relies on genuinely differing effects of adult and juvenile stress, and ideally this dissociation would be replicated before firm conclusions are drawn regarding resilience effects of stress being modulated by age of experience.

Regardless, the fact that there were no interactions between stress and *Cacna1c* manipulations implies that, at least in the present case, the effects of stress were not mediated via stress effects on *Cacna1c* expression.

Returning to the general observation that *Cacna1c*^+/-^ rats display anhedonic reactions to sucrose, the presence of this reliable deficit implies that the *Cacna1c*^+/-^ rat provides a highly valuable test bed for investigating the mechanisms by which deficiencies in VGCC function contribute to the presentation of an anhedonic phenotype. Given the fact that the *Cacna1c*^+/-^ rat also displays attenuated latent inhibition deficits characteristic of cognitive dysfunction observed in psychosis (Tigaret et al., 2021), as well as deficits in fear and reversal learning (A. L. Moon et al., 2020; Sykes et al., 2019) it affords the possibility of investigating the commonalities and potential differences in the role of VGCCs across cognitive and hedonic functions. In addition, the fact that the *Cacna1c*^+/-^ rat displays a reliable analogue of anhedonia, a symptom observed transdiagnostically across many of the psychiatric disorders for which variation in *CACNA1C* presents as a risk factor, suggests this rat may play a valuable role in the translational investigation of anhedonia more generally.

The fact that both some of the cognitive and synaptic plasticity deficits in the *Cacna1c*^+/-^ rat can be rescued by activation of the ERK pathway (Tigaret et al., 2021) suggests an obvious target for initial investigation of the cellular mechanisms underpinning *Cacna1c*-related hedonic deficits. Thus, drugs impacting VGCCs (e.g., Harrison et al., 2022) or downstream pathways (e.g., Indrigo et al., 2023) are of potential interest as candidate therapeutic approaches for both cognitive and hedonic deficits associated with genomic variation in CACNA1C.

## Notes

### Competing Interest Statement

The authors have declared no competing interest.

